# A modular method for rapidly prototyping targeted gas vesicle protein nanoparticles

**DOI:** 10.1101/2025.08.06.668980

**Authors:** Reid Vassallo, Bill Ling, Ernesto Criado-Hidalgo, Nicole Robinson, Erik Schrunk, Ann Liu, George Daghlian, Hongyi R Li, Margaret B Swift, Dhiraj Mannar, Dina Malounda, S Larry Goldenberg, Septimiu E Salcudean, Mikhail G Shapiro, Peter C Black, Michael E Cox

## Abstract

Gas vesicles (GVs) are air-filled protein nanoparticles which are proving to be useful in a number of biomedical applications. We hypothesized that it could be possible to develop a modular method for creating rapidly prototyped GVs by modifying their surface chemistry to include targeting peptides in an orientation-specific manner. Here, we describe a modular method to create targeted GVs using His-tagged antibody fragments, ensuring that the antibody fragments are connected to the GV in an orientation-specific manner. This is achieved via the functionalization of the GVs with nickel-nitrilotriacetic acid (Ni-NTA) group. First, we validated that these functionalized GVs can bind His-tagged green fluorescent protein and characterized the particle size and surface charge of functionalized GVs. Then, GVs targeted to prostate-specific membrane antigen (PSMA) using a minibody were validated using a knockout validation *in vitro*.

## Introduction

Gas vesicles (GVs) are air-filled protein nanoparticles naturally expressed by some species of cyanobacteria and archaea, which use these structures to control their buoyancy^1^. GVs have also been purified from these microorganisms, including *Anabaena flos-aquae*, to be used as nanoparticles with a variety of applications, including using them as ultrasound imaging contrast agents^2^, facilitating tumor imaging^3^, mechanotherapy^4^, cellular control^5^, enzyme sensing^6^ and calcium sensing^7^. GVs are roughly cylindrical in shape, with a diameter of approximately 85 nm and a length of approximately 500 nm^8^.

The main structural protein of GVs from *Anabaena flos-aquae*, the type of GV used here, is a small (<8 kDa^9^) gas vesicle protein A (GvpA). Each GV consists of approximately 38,600 of these self-assembling monomers^10^. There is an additional GV protein called gas vesicle protein C (GvpC), which confers GV shell stiffness^11^. The stripping of GvpC from the vesicles allows GVs to backscatter specifically designed non-linear ultrasound signals^12^.

The potential utility of GVs could be further expanded by efficiently and specifically targeting them to particular surface receptors on cells of interest through the attachment of targeting molecules to the GV surface. Such targeting has been demonstrated previously with the general integrin extracellular matrix ligand, RGD peptide^4,13^, This capability could be further expanded if it were possible to rapidly prototype a range of targeted GVs using various targeting molecules, with minimal need to tailor the synthesis pathway for each application.

Antibody conjugation has historically been a popular method of nanoparticle targeting, and has been further improved and expanded in scope with the use of antibody fragments^14^. These fragments are engineered to be smaller in size than full antibodies by removing parts of the protein which do not directly contribute to antigen binding. These modifications can lead to various benefits such as more favorable pharmacokinetics or the targeting of epitopes which were previously inaccessible due to steric hindrance^15^.

Previous work has focused on conjugating antibodies and antibody fragments to nanoparticles through various techniques^14^, including click chemistry approaches^16^. However, many of these methods utilize amino acid side chains that are present throughout the protein, so the resulting attachment of the antibody to the nanoparticle is not orientation-specific. Unsurprisingly, previous work has established that conjugating targeting molecules in an orientation-specific manner can result in improved targeting^17,18^. This is even more important for antibody fragments due their smaller size, meaning a higher proportion of the amino acids will be in the binding domain. In addition, the antibody fragment should be attached to the nanoparticle in such a way that the binding domain is facing outward to facilitate contact with its target in its environment.

Here, we describe a modular method to create rapidly-prototyped targeted GVs with orientation-specific attachment of antibody fragments (represented in Figure 1A). This orientation-specificity is achieved using nickel-nitrilotriacetic acid (Ni-NTA) conjugated to the GV surface using an N-hydroxysuccinimide (NHS) ester reaction, and the histidine (His) tag commonly present on recombinant proteins as a byproduct of the purification process. This builds upon previous work to create targeted nanoparticles using Ni-NTA by applying it specifically to GVs^17^. This modular system will allow for the rapid prototyping of targeted GVs with targeting antibody fragments for *in vitro* assessment.

**Figure 1.**
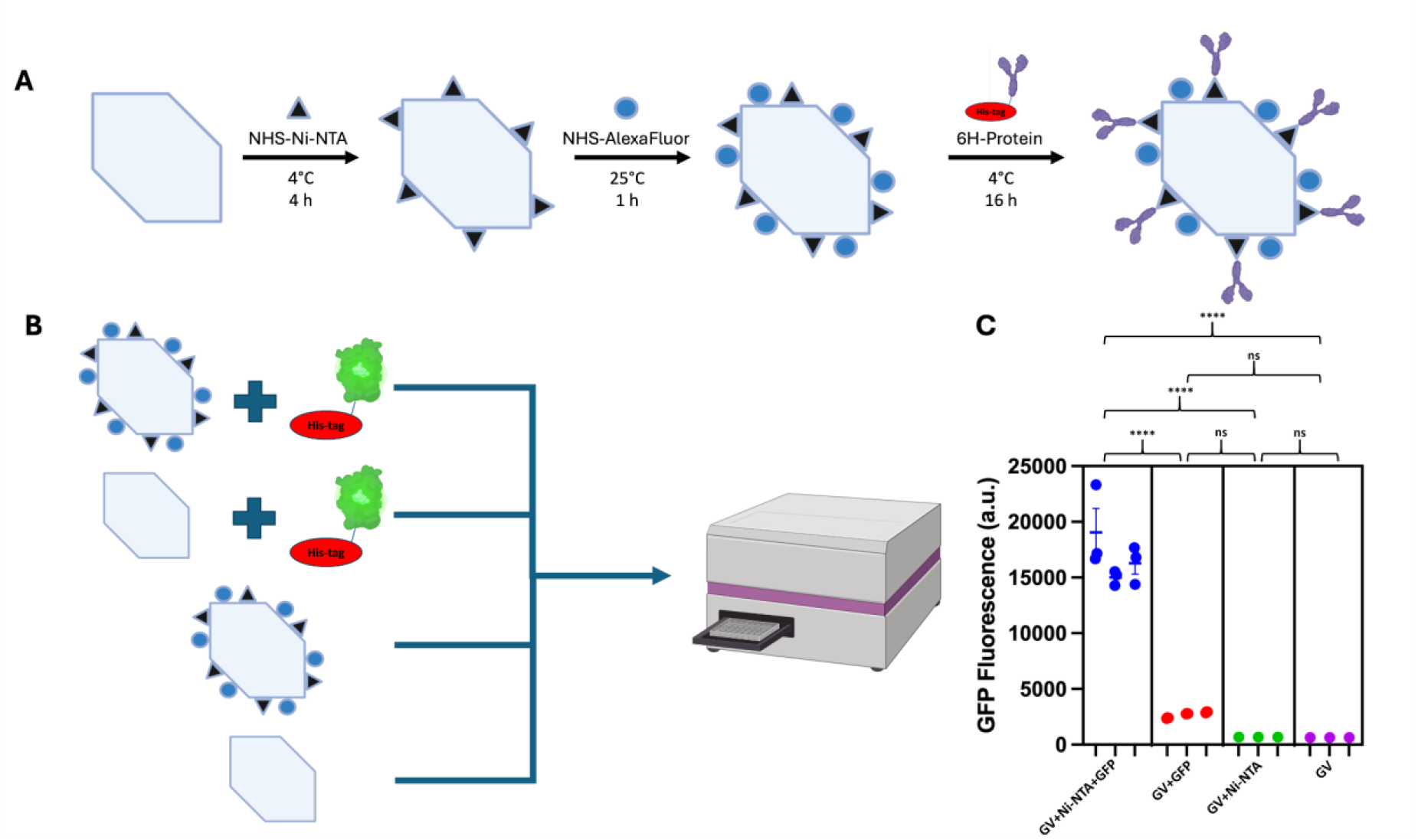
GVs functionalized with Ni-NTA are able to non-covalently bind His tag-containing proteins. (A) Representation of the functionalization process used here. For these experiments, Alexa647 is the fluorophore used. The final step shows the attachment of a generic His-tagged protein. (B) Overview of the methods of the validation experiment to assess. Fluorescent signal was compared. (C) Comparison of GFP-derived fluorescent signals in the various groups. The two binding conditions with 6H-GFP contain 3 biological replicates, each with 3 technical replicates. Conditions without 6H-GFP contain 3 technical replicates each. Statistically significant signals indicate that functionalized GVs are able to bind 6H-GFP. A lack of appreciable signal in the third condition (GV+Ni-NTA) indicates a lack of overlap in the spectrum of Alexa647 and GFP. Asterisks indicate statistical significance by a nested one-way ANOVA test with a Tukey pairwise test to correct for multiple comparisons (****: p < 0.0001, ns: not significant).

## Results and Discussion

### GVs functionalized with Ni-NTA can bind His-tagged proteins

Our modular antibody fragment-based targeting approach relies on the functionalization of GVs with Ni-NTA. The first step to validate our method of creating modular targeted GVs was to confirm that the Ni-NTA retained its ability to effectively bind His-tagged proteins when attached to the GV surface. We performed this validation using 6 His-tagged green fluorescent protein (6H-GFP) as a model protein. We expressed 6H-GFP using BL21(DE3) Competent Cells and a GFP plasmid which was under control of the endogenous T5 promoter and the lac operon.

We modified the GVs in a two-step process involving NHS ester reactions using NHS-Ni-NTA and NHS-AlexaFluor reagents, respectively, to add Ni-NTA functional groups and AlexaFluor fluorophores to the GV surface. We purified GVs from *Anabaena flos-aquae* using previously established and reported protocols^10^ and incubated these purified GVs with pre-charged NHS-Ni-NTA under gentle rotation at 4°C for 4 hours before purification by overnight dialysis against 4 L of phosphate buffered saline (PBS) and subsequent buoyancy purification^10^. We then incubated the GVs with NHS-AlexaFluor647 for 1 hour under gentle rotation at room temperature, and further purified using dialysis and buoyancy purification. The spectrum from Alexa647 does not overlap with that from GFP, which is confirmed based on the lack of fluorescence signal in the third condition in Figure 1C.

To assess whether Ni-NTA groups on the GV surface retained their ability to bind His-tagged proteins, functionalized and non-functionalized GVs were incubated with purified GFP at 4°C overnight at a ratio of 1 GFP for every GvpA protein. We determined GV concentration by quantifying their optical density at 500 nm (OD_500_) using a NanoDrop (Thermo Scientific). The amount of GvpA monomers can then be calculated using the above-mentioned ratio of 38,600 GvpA monomers per GV, considering that an OD_500_ of 1 is equivalent to 184 pmol of GVs^19^. We then used 4 rounds of buoyancy purification^10^ to remove unbound GFP in solution. The resulting GFP signal of each condition was measured using a plate reader (Figure 1B). The experiment also included 3 replicates of each of functionalized and non-functionalized GVs without 6H-GFP as negative controls. The two 6H-GFP conditions consisted of 3 biological replicates, with 3 technical replicates of each.

We observed that Ni-NTA-conjugated GVs readily bound 6H-GFP, while fluorescence intensity of non-functionalized GVs was indistinguishable from that of Ni-NTA-conjugated GVs not exposed to 6H-GFP and parental GVs alone (Figure 1C).

### Surface functionalization confirmed by differential GV particle size and surface charge

We also characterized our functionalized GVs using dynamic light scattering (DLS) to assess particle size (Figure 2A) and surface charge through zeta potential (Figure 2B) to assess whether these modifications to the GV surface significantly altered the clustering behaviour or surface properties of these nanoparticles. We compared these measurements for native GVs without GvpC, and Ni-NTA-functionalized GVs with either His-tagged human serum album (HSA) or a His-tagged minibody against prostate-specific membrane antigen (PSMA)^20^. HSA was used as a control model protein in these experiments, while the anti-PSMA minibody was chosen as a targeted GV construct. These are called HSA-GVs and PSMA-GVs, respectively, and do not contain GvpC. HSA-GVs and PSMA-GVs were created by incubating Ni-NTA-functionalized GVs with either His-tagged HSA or a His-tagged anti-PSMA minibody overnight at 4°C under gentle rotation, followed by four rounds of buoyancy purification.

**Figure 2.**
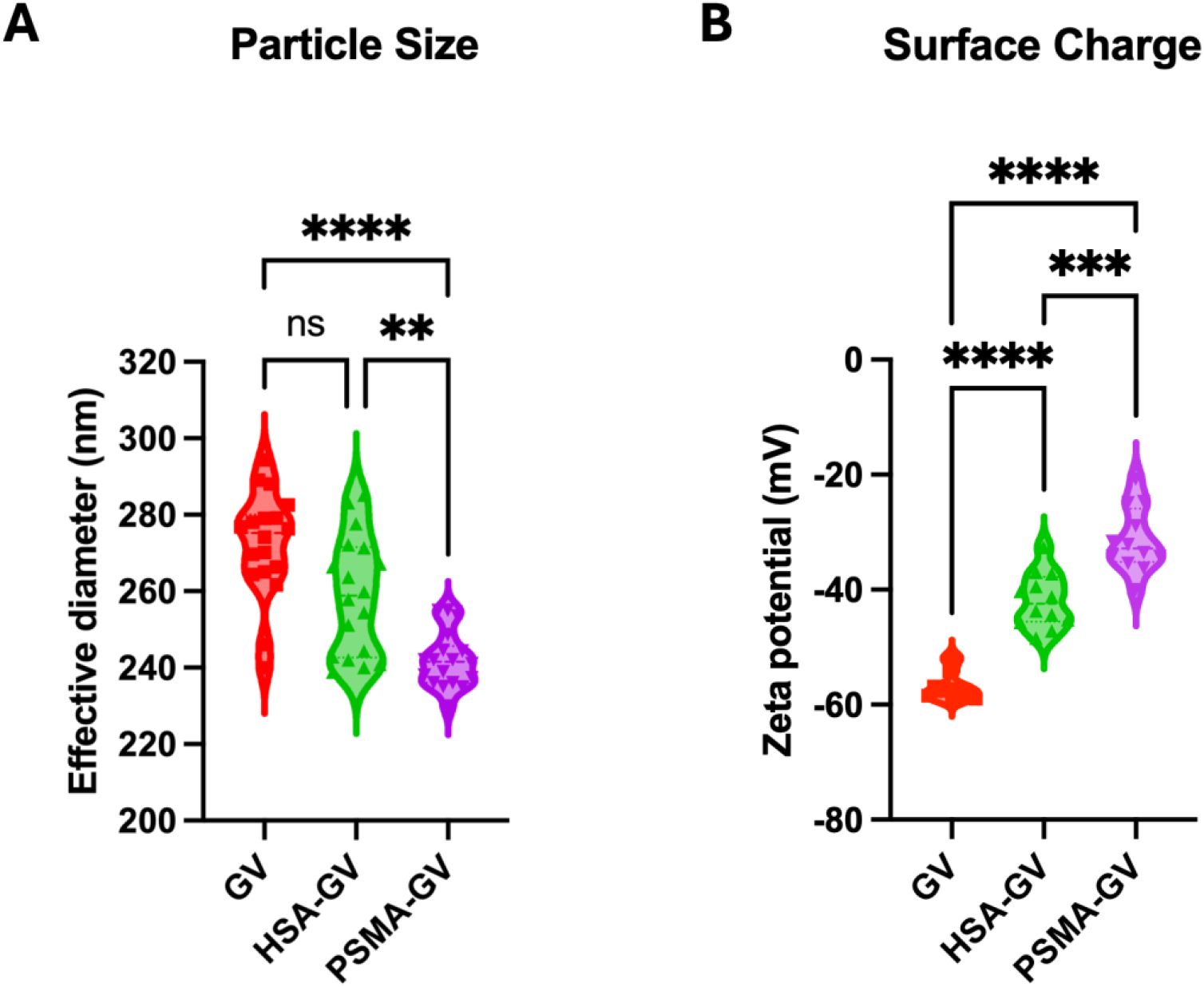
Functionalizing GVs induces minor changes in particle size and surface charge. Violin plot of (A) particle size measurements of various GV types acquired using DLS, and (B) zeta potential measurements of various GV types. From left to right in each panel, they are: native GVs without GvpC, Ni-NTA-functionalized GVs with HSA, and Ni-NTA-functionalized GVs with a minibody against PSMA. Asterisks indicate statistical significance by Welch”s t-test (****: p < 0.0001, ***: p < 0.001, **: p < 0.01, ns: not significant).

Regarding particle size, we expect HSA-GVs and PSMA-GVs to be slightly different than GVs without GvpC due to the attachment of additional protein to the GV shell, which would likely impact any slight clustering behaviour of the GVs.

The surface charges reported here for GVs without GvpC largely agree with those previously reported in the literature^21^. We see modest differences in surface charge with the addition of HSA or a minibody. These small changes are expected because both proteins have heterogeneous surface charges^22,23^.

These minor changes in our experimental measurements, namely a change in particle size and variability in the surface charge, are consistent with the expected changes, validating that modifications to the GV surface are in fact occurring.

### Targeted GVs bind preferentially to PSMA-expressing cells in a knockout validation

We assessed the targeting ability of our nanoparticles after the attachment of a minibody specific to prostate-specific membrane antigen (PSMA), a well-established cell surface biomarker for prostate cancer. We used a previously-described anti-PSMA minibody^20^ and selected the 22RV1^24^ prostate cancer cell line as our model system due to its endogenous expression of PSMA (Figure 3). To facilitate validation of PSMA-specific targeting, we utilized a PSMA knockout (PSMA KO) 22RV1 cell line, gifted by Dr. Nathan Lack of UBC, to perform a knockout validation.

**Figure 3.**
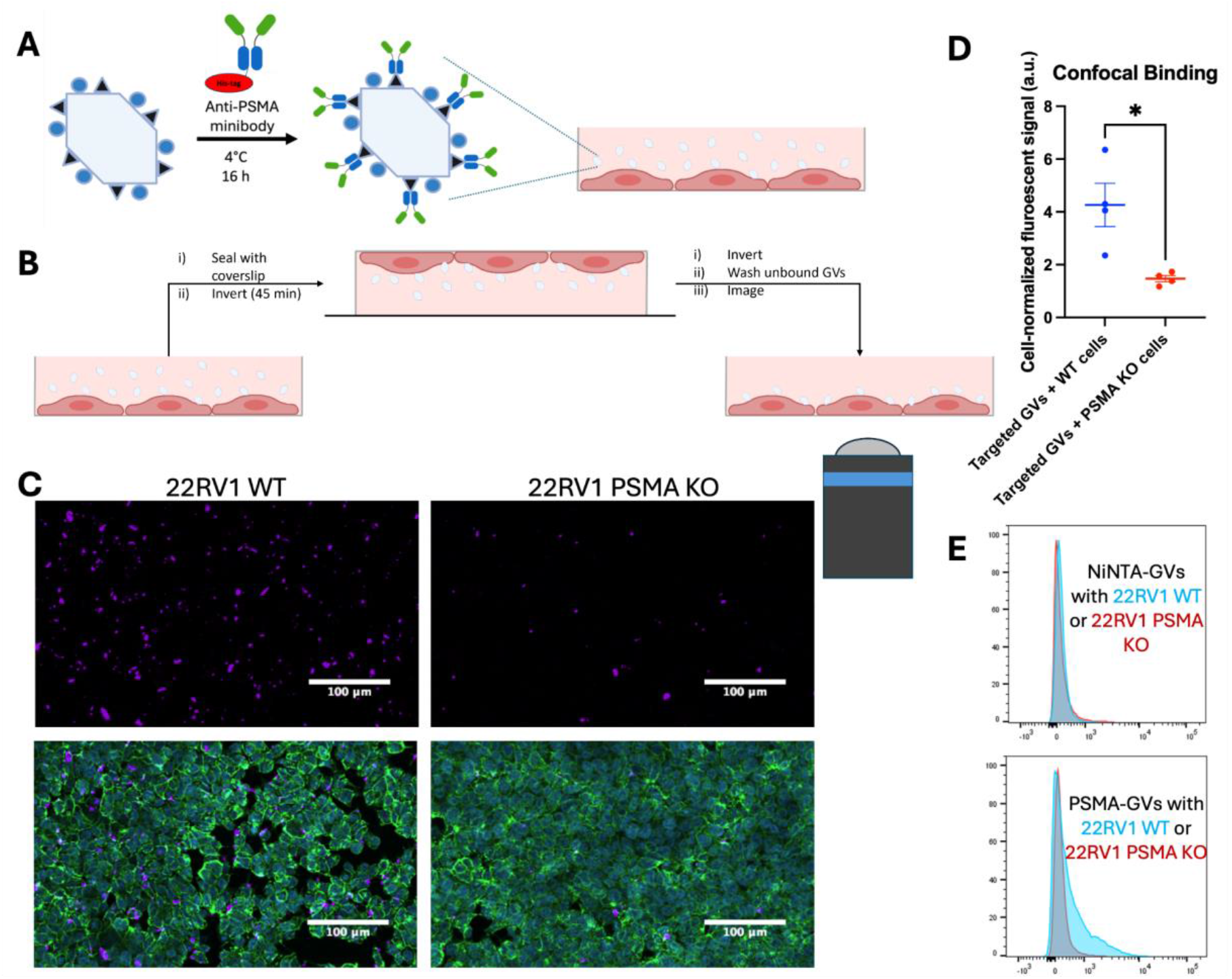
Minibody-functionalized GVs bind preferentially to cells expressing the minibody target protein. (A) Representation of the method used to attach anti-PSMA minibody to the GV surface. (B) Simplified representation of the incubation methods used in the knockout validation experiment. (C) Representative confocal images of results of the knockout validation. The left column is PMSA-expressing 22RV1 PCa cells, while the right column is PSMA-knockout 22RV1 cells. Top images are only the GV fluorescence channel, while the bottom row also contains a cell membrane stain (wheat germ agglutinin (WGA); green) and cell nuclei stain (DAPI; blue). (D) Quantification of the cell-normalized fluorescent signal between the two conditions across the entire cell samples. Asterisks indicate statistical significance by Welch”s t-test (*: p < 0.05). (E) Representative flow cytometry histograms from binding experiments using PSMA-expressing 22RV1 cells or PSMA-knockout 22RV1 cells and GVs functionalized with Ni-NTA group alone or those with the anti-PSMA minibody. This demonstrates the differential binding of the targeted GVs, as well as showing non-targeted GVs do not show preferential binding to 22RV1 WT cells.

We created targeted GVs using the above-mentioned Ni-NTA functionalization steps, followed by overnight incubation at 4°C under gentle rotation with the His-tagged anti-PSMA minibody (Figure 3A). Four rounds of buoyancy purification were performed to remove remaining free minibody in solution.

We used the same batch of targeted GVs for both conditions in this experiment to control for batch effects. WT and PSMA KO 22RV1 cells were seeded onto glass-bottom dishes and incubated overnight. The next day, they underwent a blocking step with bovine serum albumin before a 1 hour incubation with the targeted GVs. Due to GVs” buoyancy, cell cultures were partially sealed with a coverslip and inverted for the incubation period to allow for adequate contact between the cells and the GVs, following a previously published protocol^25^. Unbound GVs were removed with 6 washing steps (5 with warm medium and 1 with PBS) prior to cell staining and fixation. Confocal imaging was performed using a Zeiss LSM 800 confocal microscope in ZEN Blue (Figure 3B).

Representative images from these samples qualitatively show increased GV signal in the 22RV1 WT condition compared to 22RV1 PSMA KO, despite the similar amount of cell density (Figure 3C). We then quantified fluorescence signals from bound GVs across the entire dish in each condition and normalized against cell density, as determined by DAPI signal intensity (Figure 3D).

We further validated specific binding of targeted GVs to PSMA-expressing cells in suspension using flow cytometry. GVs functionalized with a fluorophore and Ni-NTA group (as in Figure 1A) or PSMA-GVs were incubated for 1 hour with either WT 22RV1 or PSMA KO 22RV1 cells, and fluorescence was measured using flow cytometry. As expected, both WT and PSMA KO cells incubated with NiNTA-GVs did not show any fluorescence to indicate GV-binding, demonstrating that in the absence of the anti-PSMA minibody, the Ni-NTA group itself has no binding capacity to PSMA on the cell surface. However, after incubation with PSMA-GVs, WT 22RV1 cells showed a significant increase in fluorescence compared to PSMA KO cells (Figure 3E).

These results indicate that GVs functionalized with Ni-NTA are a suitable backbone to create targeted GVs *in vitro*, but additional work will be needed to determine whether prototype targeted nanoparticles made using this method can be effectively used for *in vivo* applications of this technology.

## Conclusions

In summary, this work establishes a platform for the rapid prototyping of targeted GV nanoparticles that allows for the attachment of targeting protein fragments in an orientation-specific manner. We showed that Ni-NTA retains its ability to bind His-tagged proteins when it is used to functionalize the GV surface, and induces minor changes to the effective particle diameter and surface charge. It was then demonstrated using an *in vitro* knockout validation that targeted GVs produced using this method preferentially bind to PSMA-expressing cells. These methods can be further used to create targeted nanoparticle prototypes with targeting protein fragments for rapid *in vitro* assessment of their targeting ability. Additional work is needed to assess if this system is reliable to assess *in vivo* targeting ability.

## Experimental Methods

### GV Functionalization

We functionalized GVs in a modular fashion using N-hydroxysuccinimide (NHS) ester reactions and nickel-nitrilotriacetic acid (Ni-NTA). This was done to add a previously-described PSMA-targeted minibody^20^ and a fluorophore to the GVs for validation purposes.

Ni-NTA was used to attach the minibody to the GV surface in order to preserve the proper orientation of the minibody where the binding domain is facing away from the GV surface^17^.

#### Creation of pre-loaded NHS-NiNTA

We pre-loaded NHS ester-NTA (NHS-NTA) (AAT Bioquest, Pleasanton, CA, USA) with Ni^2+^ to avoid aggregation and maximize binding^17^. To do this, NHS-NTA was aliquoted into 100 mM stock solution in dimethyl sulfoxide (DMSO) for long-term storage at −20°C. After thawing and immediately before use to functionalize the GVs, the NHS ester-NTA solution was added to a 10-fold excess volume of a freshly prepared 0.5 M solution of nickel chloride hexahydrate (NiCl_2_-6H_2_O) in borate buffered saline (BBS) for 5 minutes at room temperature, resulting in NHS-NiNTA.

#### Construction of PSMA-GVs

Stripped Anabaena GVs were produced and purified following previously published protocols^10^. To ensure adequate functionalization, we reacted GVs with a 10-fold molar excess of NHS-NiNTA per GvpA, the primary structural protein of the GV^26^. This corresponds to a 386,000-fold molar excess of NHS-NiNTA per GV particle^10^. The reaction was allowed to proceed under gentle rocking for 4 hours at 4°C. Purification of particles was performed by first dialyzing against 4 L of PBS overnight, followed by a round of buoyancy purification. Buoyancy purification of stripped Ana GVs was performed by centrifugation at 300 x g for 3 hours. Another NHS ester reaction was performed to label the GVs with an NHS-Alexa fluorophore (Thermo Fisher, Waltham, MA, USA), at 10 000-fold molar excess. For most experiments Alexa 647 was used, but based on instrument differences Alexa 488 was used for the flow cytometry experiment in Figure 3E. This reaction was again followed by dialysis and buoyancy purification.

A previously-described anti-PSMA minibody^20^, known to have a 6-histidine tag at the non-binding C-terminal end, was then added to the functionalized GVs at a ratio of 1 minibody per 60 gvpAs. For particle characterization experiments, we also followed this step with the same molar amount of His-tagged human serum albumin (HSA) (AcroBiosystems, Newark, DE, USA).

### Green fluorescent protein production

We used a methionine-less GFP variant called GFP-1M which is under the control of the endogenous T5 promoter and the lac operon. The protein was produced using BL21(DE3) Competent Cells. The BL21(DE3) cells were transformed using the GFP-expressing plasmid which contained an ampicillin-resistance gene. After transformation, cells were cultured overnight in 37°C on a plate containing Luria Broth (LB) (Invitrogen, Waltham, MA, USA), 1.5% weight/volume (w/v) agar, 1% w/v glucose, and 200 μg/mL ampicillin. This step was selected for cells which had successfully taken up the GFP-expression plasmid. Then, a culture was again grown overnight with two colonies picked from the plate in 3 mL of LB with 1% w/v glucose and 200 μg/mL ampicillin. After growing overnight, this 3 mL saturated culture was split into two 250 mL Erlenmeyer flasks containing 100 mL LB with 0.2% w/v glucose and 200 μg/mL ampicillin. They were each induced with 1 mL of 100 mM isopropyl β-D-1-thiogalactopyranoside (IPTG) after 3 hours and allowed to grow for a further 6 hours, all at 37°C. Following this, the cultures were centrifuged at 5000 x g for 10 minutes and the cell pellets were kept. They were soaked in 9 mL of Solulyse™ Bacterial Protein Extraction Reagent (AMS Biotechnology, Oxfordshire, UK) overnight for cell lysis. GFP was then retrieved using a Ni-NTA resin column. After elution with imidazole, the GFP underwent 2 rounds of overnight dialysis against 4 L of PBS to remove the excess imidazole so it would not interfere with future steps. GFP concentration was determined using a Pierce™ 660nm Protein Assay (Thermo Scientific, Waltham, MA, USA).

### Particle characterization

We characterized modified GVs using dynamic light scattering (DLS) to study their size, and zeta potential to measure their surface charge. Here, native GVs without GvpC were compared against Ni-NTA-functionalized GVs coated with either HSA or a PSMA-targeting minibody. The DLS and zeta potential procedures largely follow those previously used by Ling *et al*.^13^, but they are described here as well. To compare between groups, an analysis of variance (ANOVA) test was used, followed by pair-wise Welch”s t-tests with a Dunnett”s correction for multiple comparisons.

#### Particle size measurements

A GV suspension of 25 uL OD_500_ 2 GVs in 500 uL PBS was dispensed into a disposable semi-micro polystyrene cuvette (VWR, Radnor, PA, USA). This was then measured using a ZetaPALS particle analyzer (Brookhaven Instruments, Nashua, NH, USA). Particle size was measured by the ZetaPALS Particle Sizing software using an angle of 90°, the thin shell setting, a run length of 15 seconds, and 6 runs per sample. Measurements were recorded as mean intensity-weighted diameter. This was performed on 3 replicates (18 measurements total).

#### Surface charge measurements

An electrode (SZP, Brookhaven Instruments, Nashua, NH, USA) was inserted into a mixture of 50 uL OD 2 GVs in 1.5 mL Milli-Q water in a disposable plastic cuvette (SCP, Brookhaven Instruments). Measurements were performed using the ZetaPALS Zeta Potential software using the Smoluchowski model, based on 4 measurements of 15 cycles each. This was performed on 3 replicates (12 measurements total).

### Binding Experiments

For all mammalian cell culture, the media used was Roswell Park Memorial Institute (RPMI) 1640 media (Corning Life Sciences, Corning, NY, USA), supplemented with 10% v/v fetal bovine serum (FBS), 1% v/v GlutaMAX (Thermo Fisher Scientific, Waltham, MA, USA), and 2% v/v Antibiotic-Antimycotic (Gibco, Waltham, MA, USA). A well-established PSMA-positive cell line, 22RV1^24^, was used for these experiments. Wild type (WT) 22RV1 cells were used as the PSMA-positive condition, while a PSMA-knockout (PSMA KO) version of this cell line was used as our PSMA-negative condition. The PSMA KO cell line was a gift from Dr. Nathan Lack at the University of British Columbia.

#### Confocal microscopy experiment

The ability of PSMA-GVs to bind to PSMA-positive PCa cells *in vitro* was validated using confocal microscopy. For these experiments, approximately 200,000 WT or PSMA-knockout 22RV1^24^ cells were cultured on fibronectin-coated glass-bottomed 35 mm cell culture dishes (Matsunami Glass, Kishiwada, Japan) overnight. Then, these cells were incubated with 200 μL of a blocking solution (2% w/v bovine serum albumin (BSA) in PBS) for 45 minutes at 37°C. Following this, they were incubated in RPMI media for 15 minutes for recovery. Then, they were incubated with 200 μL of PSMA-GVs at an OD_500_ value of 1 for 1 hour at 37°C while inverted. Following previously reported methods^25^, the cells are inverted at this step so the cells remain in contact with the buoyant GVs. The cells were then washed five times with media and once with PBS. The cells were then incubated with 5 μg/mL wheat germ agglutinin (WGA) (conjugated to Alexa Fluor 488 (Invitrogen, Waltham, MA, USA)) to stain for cell membranes for 20 minutes, followed by fixation using 4% paraformaldehyde, and staining with 4”,6-diamidino-2-phenylindole (DAPI) at a 5 μg/mL concentration, to visualize the cell nuclei.

#### Flow cytometry experiment

22RV1 cell cultures (WT and PSMA KO) were incubated with 0.25% Trypsin-EDTA (Gibco, Waltham, MA, USA) until lifted. Once complete, this reaction was stopped with RPMI medium supplemented with 10% FBS (Gibco, Waltham, MA, USA). Cells were resuspended in fluorescence-activated cell sorting (FACS) buffer (120 mM NaCl, 5 mM KCl, 5 mM MgCl_2_, 40 mM HEPES, pH 7.5, 0.01% BSA, 0.15% glucose) before being counted using trypan blue and a TC 20 automated cell counter (Bio-Rad, Hercules, CA, USA). After counting, the cell suspension was separated so as to have 200,000 live cells per sample tube. Cells were incubated for 60 minutes in FACS buffer containing GVs with an OD_500_ value of 2 and a LIVE/DEAD™ Fixable Dead Cell Stain at room temperature on a rocking platform. Following this, cells were washed 3 times in FACS buffer and data was collected on a FACS Canto II (BD Biosciences, Franklin Lakes, NJ, USA). Analysis was performed using FlowJo software (FlowJo, Ashland, OR, USA).

### Cell imaging and image analysis

Confocal imaging was performed using a Zeiss LSM 800 confocal microscope in ZEN Blue (Carl Zeiss AG, Oberkochen, Germany), and image analysis was performed using MATLAB 2024a (The Mathworks, Matick, MA, USA).

To acquire representative qualitative images, we acquired 6×6 tiled images using a 20x objective lens, resulting in a field of view of 1.8 mm x 1.8 mm. In order to avoid unintended sampling bias, to quantify the binding we acquired tiled images of the entire glass-bottom dish using a 10x objective.

For these quantification images, the Alexa 647 signal (corresponding to the amount of GVs present) was calculated, and was normalized to the amount of cells present using the DAPI signal. First, the round glass-bottomed dish was manually segmented in order to avoid artifacts around the edge of the glass-bottomed section. Then, determination of the relative amount of cells present was done using a threshold-based segmentation approach, where pixels with a value of at least 400 were deemed to include a cell nucleus. The total number of these pixels in the image were summed, and the Alexa 647 signal was divided by this number.

The cell-normalized fluorescence signal was then compared between WT and PSMA KO cell conditions using a Welch”s t-test to account for potentially unequal variances.

## Acknowledgements

We would like to thank the following collaborators for helpful discussions throughout this work: Rohit Singla, Nivin Nystrom, Di Wu, Yuxing Yao, Shirin Shivaei, Rohit Nayak, Mitali Pandey, Hao Shen, Nathan Lack, Flora Huang, Meric Dikbas, and Atsuhiko Yoshizawa.

Funding for this work was provided by the Friedman Award for Scholars in Health from the University of British Columbia for R.V., a Natural Sciences and Engineering Research Council (NSERC) Canada Graduate Scholarship and Michael Smith Foreign Study Supplement for R.V., the Lazlo Chair in Biomedical Engineering for S.E.S. and the National Institutes of Health (RO1-EB018975) for M.G.S.

